# A novel lipase with dual localisation in *Trypanosoma brucei*

**DOI:** 10.1101/2022.02.23.481587

**Authors:** S Monic, A Lamy, M Thonnus, T Bizarra-Rebelo, F Bringaud, TK Smith, LM Figueiredo, L Rivière

## Abstract

Phospholipases are esterases involved in lipid catabolism. In pathogenic micro-organisms (bacteria, fungi, parasites) they often play a critical role in virulence and pathogenicity. A few phospholipases (PL) have been characterised so far at the gene and protein level in unicellular parasites including African trypanosomes (AT). They could play a role in different processes such as host-pathogen interaction, antigenic variation, intermediary metabolism. By mining the genome database of AT we found putative new phospholipase candidate genes and here we provided biochemical evidence that one of these has lypolytic activity. This protein has a unique non-canonical glycosome targeting signal responsible for its dual localisation in the cytosol and the peroxysomes-related organelles named glycosomes. We also show that this new phospholipase is excreted by these pathogens and that antibodies directed against this protein are generated during an experimental infection with *T. brucei gambiense*, a subspecies responsible for infection in humans. This feature makes this protein a possible tool for diagnosis.

## Introduction

Trypanosomatids are protozoa transmitted by insect vectors and cause human and animal diseases. The three main species are American trypanosomes (*Trypanosoma cruzi*, Chagas disease), *Leishmania spp* (cutaneous, mucocutaneous and visceral leishmaniasis) and African trypanosomes (*Trypanosoma brucei*, sleeping sickness). Millions people are affected by these diseases and more than half a billion are at risk (World Health Organization). From a veterinary point of view, some trypanosomes species are particularly virulent on farm animals, especially in Africa where this leads to heavy economic losses and it constitutes a major obstacle to the development of this continent^1,2^.

Phospholipases (PLs) and *lyso*-phospholipases (LysoPLA) belong to a complex group of enzymes that cleave phospholipids and *lyso*-phospholipids. PLs are classified as A1, A2, B, C or D and LysoPLAs as LysoPLA1 and LysoPLA2 depending on the site of hydrolysis^3^. Numerous LysoPLA have PLA1 activity in addition of their true LysoPLA activity^3^.

These lipases have a variety of biological functions including production of bioactive lipids that act as second messengers and modulators of the immune response^4,5^. Some PLs are involved in the recycling of membrane phospholipids and others have cytolytic effects. Noteworthy, PLs are powerful toxins found in the venom of bees and snakes and trigger blood and necrotic damage^6,7^.

In several pathogens (*Pseudomonas, Ricktesia, Candida, Amoeba, Giardia, Toxoplasma* and others) phospholipases play a role in the infection and have been recognised as true virulence and pathogenic factors^8,9^.

Phospholipases A1 (PLA1) activities are capable of hydrolyzing the sn-1 acyl ester function of the phospholipids. In protozoan parasites, only a few genes encoding PLA1 have been cloned and studied so far^10,11^. In higher eukaryotes, classification of PLA1s does not rely on sequence similarity, but rather on subcellular localisation. Indeed PLA1s can be divided into two groups, the first group includes excreted enzymes, while the second one contains intracellular enzymes^10,12^.

Based on the literature and genomic data, kinetoplastids possess several proteins with putative PLA1 activity among which a few have been already experimentally described. In *T. cruzi*, a parasite with a predominantly intracellular lifestyle, the only known PLA1 is a membrane-bound and excreted protein that alters the lipid profile of the host *via* second messenger production and concomitant activation of protein kinase C^13^. *Leishmania spp* possess also a PLA1 that could be involved in virulence^14^. It has been demonstrated that the pathogens *T. brucei* and *Trypanosoma congolense* possess a higher PLA1 activity than the non-pathogenic *Trypanosoma lewisi*^15^. Moreover, in *T. brucei* mammalian forms (bloodstream form-BSF) the PLA1 activity is higher than in insect forms^16^. Finally, a strong PLA1 activity has been measured specifically in tissue fluids of *T. brucei*-infected rabbits and this activity positively varies according to the waves of parasitemia^17^. Altogether this suggests that PLA1 could play an important role in the host-pathogen interaction. In *T. brucei* no PLA2s have been described so far and it is the same for LysoPLAs.

So far, only one phospholipase gene has been described and studied in *T. brucei*. It encodes a cytosolic protein with PLA1 activity, which is involved in the synthesis of *lyso*-phosphatidylcholine (*lyso*-PC) metabolites^18^. This protein was named TbPLA1 by the authors. TbPLA1 is not excreted and is neither essential for *in vitro* growth of both parasite stages (Insect and bloodstream stages), nor for *in vivo* virulence^18,19^. Although the trypanosome genome contains a number of genes that could encode putative phospholipases (our unpublished work), those responsible for PLA1 activity are unknown.

In this study we describe some features of a novel *T. brucei* lipase already annoted as LysoPLA based on automated sequence homology analysis. We show that this lipase harbors a PLA1 activity on non-natural phospholipids and a PLA2 on natural phospholipid *in vitro*. We show also that this protein is not only distributed in both glycosomes and cytosol but is also excreted in the medium. We discuss the potential roles played by PLA in trypanosomes and its interaction with the host.

## Results

### In silico identification and sequence analysis of a new putative lipase in Trypanosoma brucei

Previous studies strongly suggested the existence of more than one phospholipase in African trypanosomes playing important role(s) in host-pathogen interaction^10,11^. Our goal was to identify such new phospholipase(s) in *T. brucei*. We rationalised our research by combining an *in silico* analysis with a literature analysis, including global approaches described below. We found a gene, Tb927.8.6390 (TriTrypDB accession number), with a predicted α/β hydrolase / phospholipase domain^20^ and annotated as TbLysoPLA (standing for *Trypanosoma brucei* Lyso-phospholipase A) based on sequences homology.

According to the literature^21,22^ this gene encodes for a protein located in the glycosomes which are peroxisome-like organelles^23^ and also is possibly excreted/secreted^24^. Moreover, based on the previously published RNAi screen (performed by Alsford et al^25^), the gene seems to be essential in BSF. Orthologs of Tb927.8.6390 in other kinetoplastids have a very good general conservation of the amino acid sequence (for example 53% identities with *leishmania* and 70% identities with *T.congolense*) (Figure 1A and table S1) in which two elements are easily recognisable. First, the putative active site typical of *lyso*-phospholipases and phospholipases (GXSXG^26^) has been identified (Figure S1) and is perfectly conserved in all these organisms (Figure 1, box1), confirming that the protein should have an hydrolase activity. Then, the carboxy-terminal end is not conserved (Figure 1, box2). For a protein to be imported into the glycosomes, it must have a specific targeting signal such as the canonical Peroxisome-Targeting Signal 1 (PTS1) which is a tripeptide (S/A/C)(K/R/H)(L/M) found in the C-terminus^27^. As mentioned above, Tb927.8.6390 has been detected in glycosomes in several mass-spectrometry analysis^21,22^ and this localisation could be due to the tripeptide “SKS” at the extreme C-terminus. This sequence does not correspond strictly to a canonical PTS1 and has not been observed as a PTS1 in other kinetoplastid species. Since the “SKS” is unusual, we searched in the TriTryp genomic database for other *T. brucei* encoded proteins that possess the same motif. The table presented in Figure 1B shows that two other proteins contain this C-terminal tripeptide. The transmembrane adaptator Erv26 is found in COPII vesicles^28^ and CIF1 is located at the cell tip^29^.

**Figure 1:**
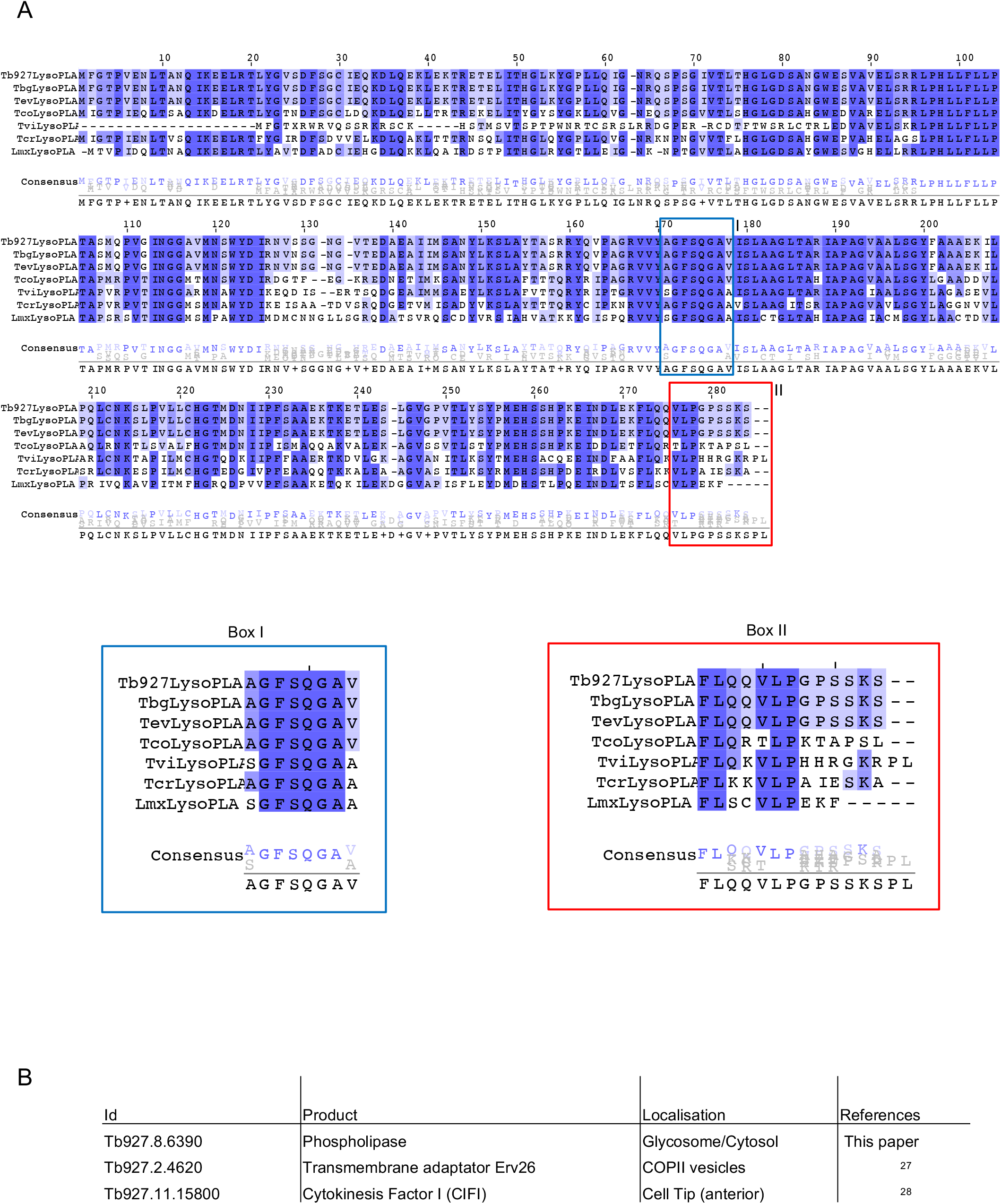
Comparison of LysoPLA in kinetoplastids. A / Alignment of LysoPLA from kinetoplastids. Sequences were extracted and aligned using Clustal omega. Dark blue contains conserved residues, white to light blue contains conservative changes. Box I emphasizes the conservation of the putative phospholipase active site. Box II emphasizes the differences between kinetoplastids LysoPLA Carboxy-terminus. Tb, *Trypanosoma brucei brucei (Tb927.8.6390*); Tbg, *Trypanosoma brucei gambiense (Tbg972.8.6450*), Tev, *Trypanosoma evansi* (TevSTIB805.5.6680); Tco, *Trypanosoma congolense (TcIL3000_8_6240*); Tv, *Trypanosoma vivax (TvY486_0805980*); Tcr, *Trypanosoma cruzi (TcCLB.506797.70*); Lmx, *Leishmania Mexicana (LmxM.24.1840*). B / Only 3 proteins are ending with SKS in *Trypanosoma brucei brucei*. The search was performed on TriTrypDB.org using the « Protein Motif Pattern » tool.

### Tb927.8.6390 encodes for a protein with lipolytic activity in vitro

As said before, based on sequence homology, Tb927.8.6390 has been annotated as a *lyso*-phospholipase. These enzymes have a broad spectrum of lipolytic activities including phospholipase A. Instead of dissecting the precise activity of this protein, we wanted to proof that this enzyme displays a lipolytic activity. We decided initially to test this by using commercial *in vitro* phospholipase A detection kits. No *lyso*-phospholipase assay kits are commercially available. The purified recombinant protein expressed in *E. coli* (Figure 2A, left and S2) was tested for phospholipase activities using an *in vitro* non-natural fluorogenic assay (also see material and methods). In these assays, substrates have a cleavable fluorophore attached to either *sn-1 or sn-2* position. PLA1 activity was observed with the Fluorogenic PED-A1 (phosphatidylethanolamine with NBD-labeled only on the acyl linked *sn-1* position, while the *sn-2* position has a non-cleavable alkyl (Figure 2A, right). However, no PLA2 activity was observed with, the BODIPY-labelled phosphatidylcholine with the cleavable BODIPY in the *sn-2* position, and a non-cleavable amide linked BODIPY in the *sn-1* position (data not shown). Thus, the recombinant TbLysoPLA, has lipase activity, being able to cleave the non-natural PE *sn-1* acyl next to the *sn-2* alkyl.

**Figure 2:**
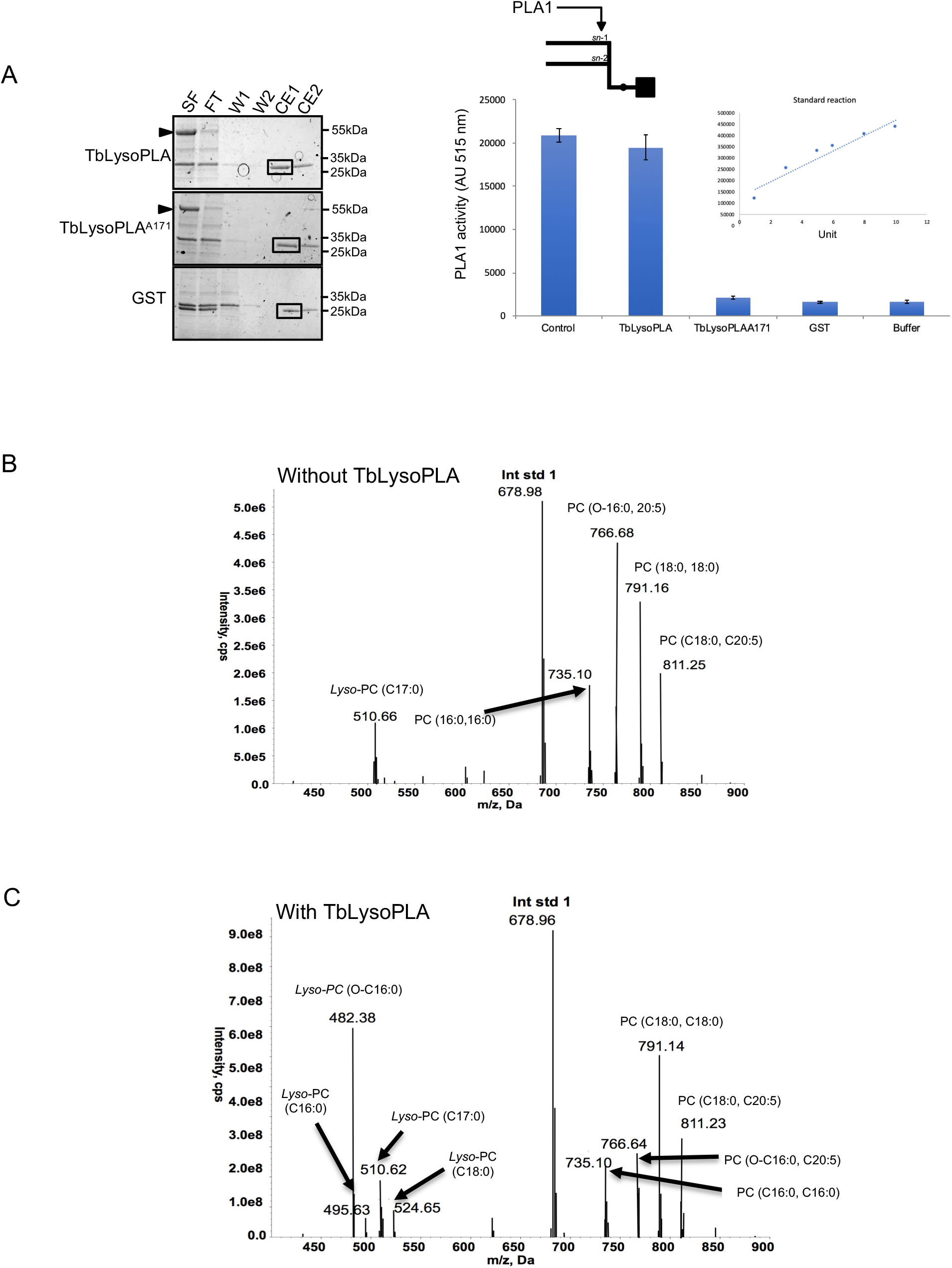
Lipolytic activity of recombinant TbLysoPLA. A / PLA1 activity assay on recombinant proteins expressed in *E. coli*. Purification steps were analysed by coomassie gel (above gel). As examplified for TbLysoPLA, SF (Soluble Fraction), FT (Flow Through), W1-2 (Washes), CE1-2 (Clivage/Elution after thrombin release). Tested fractions were dialysed against PBS. PLA1 activity was measured using Enzcheck PLA1 assay (Molecular Probes). Control was a commercial phospholipase A1 (Sigma L3295). TbLysoPLA, full-length TbLysoPLA; TbLysoPLA^SA171^, mutated TbLysoPLA where the putative active serine 171 was replaced by an alanine; GST, Glutathione-S-Transferase was eluted with 20mM glutathione instead of thrombin cleavage. Supplemental information concerning expression and purification can be found on figure S2. B/ Substrate specificity of recombinant TbLysoPLA. TbLysoPLA was incubated (B) or not (A) with a Lipid Mix containing *lyso*-PC C17:0, PC (diC16:0), PC O-C16, 20:5, PC (diC18:0).

We have also expressed and purified a mutated version, in which the putative catalytic serine (S^171^) has been replaced by an alanine (TbLysoPLA^S171A^). As expected we could not detect any phospholipase A1 activity with this S171A mutated recombinant enzyme (Figure 2A, right). Then we would like to evaluate the capability of our recombinant enzyme to cleave natural phospholipids or *lyso*-phospholipids. For that, several lipids mixed in micelles were incubated without or with the recombinant enzyme and analysed by mass spectrometry (Fig 2 B and C respectively)^19^. The lipid intensities were normalized to the internal control (PC 14:0, 14:0). and upon treatment with TbLysoPLA the formation of *lyso*-PC (16:0), *lyso*-PC (18:0) and *lyso*-PC (O-16:0) are observed, with the latter being the most dominant *lyso*-species. This strongly suggests that TbLysoPLA is able to cleave the *sn-2* C20:5 fatty acid from PC (O-16:0, 20:5). This is somewhat surprising given the fluorogenic assays with the non-natural, as described above. However, cleavage of the acyl chain next to an alkyl linkage is common to both. Additionaly, some formation of *lyso*-PC (16:0), which has come from PC (16:0, 16:0) and *lyso*-PC (18:0) again from either PC (18:0, 18:0) and /or PC (18:0,.20:5). Based upon these observations it is not possible to conclude at this stage if the cleavage occurs at the *sn-1* or *sn-2* position. The amount of *lyso*-PC (17:0), which was also presented as a possible substrate, does not decrease significantly relative to the internal standard and therefore, it is clear that TbLysoPLA, does not cleave the sn-1 acyl group from *lyso*-PC species.

Finally, the enzyme was not able to cleave natural PG, PA, PS or PI (supplemental figure S3), suggesting a specificity towards the head group of phospholipids.

Altogether these results demonstrate that the recombinant enzyme can display a PLA2 activity *in vitro*. Tb927.8.6390 is therefore a lipase. A further thorough investigation of its enzymatic specificities *in vitro* and *in vivo* is required to fully understand the catalytic properties of this protein. Nevertheless, given actual annotations we have chosen to keep the actual name TbLysoPLA.

### TbLysoPLA is constitutively expressed

The purified recombinant TbLysoPLA described before was used to immunize Rabbits in order to raise a serum directed against it (See materials and methods). This immune serum recognises a single 30 kDa protein corresponding to the size predicted in Tritryp database (Figure 3A, left). This signal is detected in both Bloodstream and procyclic cultivated stages indicating that this enzyme is constitutively expressed. As observed in the figure only one protein could be detected by the polyclonal serum, and we could not detect any signal either in the pre-immune serum or cell-lines depleted of TbLysoPLA showing the specificity of our antibody. Previous analysis suggested that anti-TbPLA1 could cross-react with TcPLA1 even if the sequences are not well-conserved^13^. TbLysoPLA and the PLA1 from Tb, Tc and Lm are also showing a low level of amino-acids identity (cf S4 A and B). As shown in figure S4C our antibody could detect the ortholog of TbLysoPLA in *Leishmania*. As a control, anti-TbPLA1 did not give any signal with leishmania extract while detection could be possible in the cellline depleted of TbLysoPLA. Altogether these results (figure 3 and figure S4) show that our antibody is highly specific and do not cross-react with TbPLA1.

**Figure 3.**
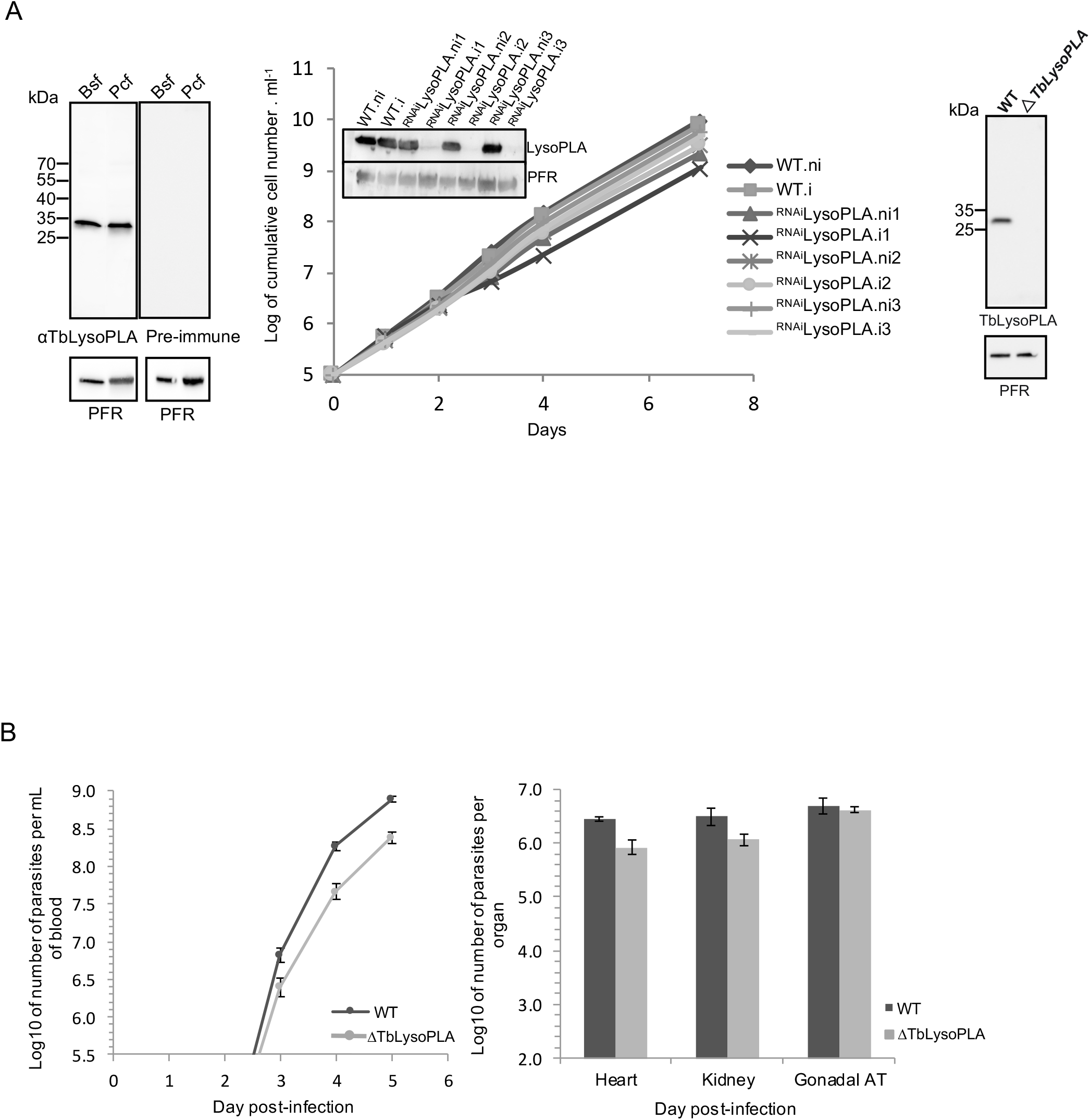
Analysis of bloodstream form mutants. A / Left: TbLysoPLA is expressed in both Bloodstream and Procyclic forms. Total protein extracts from both forms of Tb were resolved by SDS-PAGE and transferred on nitrocellulose as described in materiel and method section. Membranes were probed with sera raised against TbLysoPLA and PFR. Right panel is a control with the pre-immune sera from the animal before immunisation with TbLysoPLA. Middle: Inhibition of TbLysoPLA expression by interference RNA. Growth of the parental cellline and 3 independant clones cultivated with (i) or without (ni) tetracycline. Right panel shows the ΔTbLysoPLA validation by Western Blotting as TbLysoPLA is not detected anymore. A specific affinity-purified anti-TbLysoPLA antibody was used and PFR was used as loading control. B / Mice infection with wild-type (dark grey) and ΔTbLysoPLA (light grey) Tb427 BSF. Error bars represent the standard error of the mean (n=4 per group). left / Parasitemia of infected mice quantified by hemocytometer. right / Number of parasites in heart, kidney and gonadal adipose tissue, 5 days after infection, quantified by qPCR.

### TbLysoPLA is not essential for the survival of parasites in vitro and in vivo

According to the RITseq screen performed by Alsford et al^25^, TbLysoPLA appears to be essential for the BSF, making it a target for a possible treatment. In order to verify these results, expression of this protein was conditionally down-regulated by RNAi in the presence of tetracycline^30^. As shown in Figure 3A (middle), inhibition of LysoPLA expression in three tetracycline-induced ^RNAi^TbLysoPLA cell lines has no impact on the parasite growth *in vitro* although LysoPLA is no more detectable by Western blot (Figure 3A). Since residual amounts of the targeted protein can be expressed in RNAi cell lines, both alleles of the TbLysoPLA gene were deleted by gene replacement (ΔTbLysoPLA, Figure 3A, right panel, fig sup). The ΔTbLysoPLA mutant showed no growth defect (not shown) confirming that LysoPLA is not essential for BSF *in vitro*.

In order to test the importance of TbLysoPLA for parasite survival in the mammalian host, mice were infected with the ΔTbLysoPLA strain or the parental line. If TbLysoPLA is essential for the establishment of infection, we expected a decrease in the number of null-mutant parasites in the blood and organs. Both WT and ΔTbLysoPLA parasites show similar levels of parasitemia (Figure 3B). On day 5, parasitemia reached > 10^8^ parasites and mice were sacrificed. On this day, solid tissues were collected to count the total number of parasites by qPCR. We observed that the total number of WT parasites in the solid tissues was very high in three tissues analysed (heart, kidney and adipose tissue), which is probably a consequence of the uncontrolled parasitemia that stems from lack of pleomorphism of Lister427 strain. In any case, we observed no statistically significant differences in parasite load between WT and ΔTbLysoPLA parasites in any tissue, indicating that TbLysoPLA is not necessary for parasite survival (Figure 3B) and growth in any of the tested tissues *in vivo*.

Overall, we conclude that TbLysoPLA is not essential for the viability of the strain Lister427 neither *in vitro* nor *in vivo*.

### Tb LysoPLA localizes in the glycosomes and the cytosol

Several proteomic studies suggested that TbLysoPLA is a glycosomal protein^21,22^, which is consistent with the presence of a signal PTS1-like signal at the C-terminal extremity. We used a digitonin titration experiment to determine the localisation of the protein. In this approach the parasite membranes are differentially permeabilized using increasing concentrations of this detergent^31^. Western Blot analysis of the soluble fractions using our specific anti-TbLysoPLA shows that this protein is released together with the enolase, a cytosolic marker (Figure 4A,^32^). The analysis of the insoluble fractions shows that the profile of TbLysoPLA is similar to that of aldolase, a glycosomal marker (Figure 4A,^32^). These results indicate that TbLysoPLA is distributed in both the glycosomes and the cytosol of BSFs. This is in contrast to previously published describing LysoPLA only as a glycosomal protein^21,22^.

**Figure 4:**
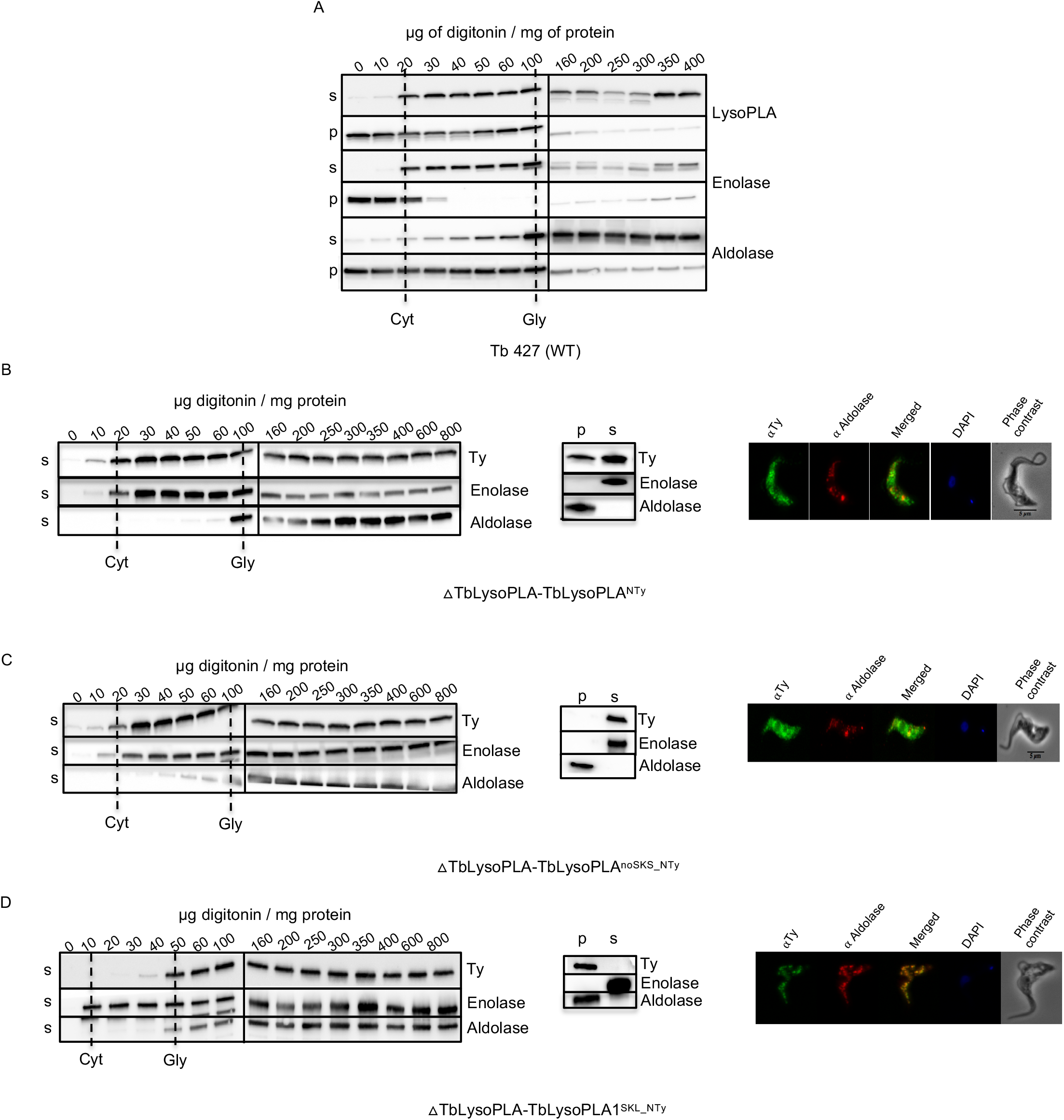
Subcellular distribution of TbLysoPLA in Bloodstream forms. Digitonin titration (left of B, C and D): Western blot analysis of the supernatant (s) and pellet (p) fractions from cells incubated with 0,01-0,4 mg digitonin/ mg protein in STE buffer containing 150 mM NaCl. The digitonin concentration required to release cytosolic (cyt) and glycosomal (gly) marker proteins are indicated by vertical dash line. TbLysoPLA was detected with specific anti-TbLysoPLA in A and anti-TY in B, C and D. For technical reasons two gels/membranes were needed to analyse the whole set of samples. The Vertical line between 100 and 160 indicates a delimitation between two different membranes. Hypotonic lysis (middle of B, C and D): Western blot analysis of supernatant (s) and pellet (p) fractions from cells incubated with hypotonic buffer. Anti-TY was used for detection of TbLysoPLA. Immunofluorescence Assay (IFA; right of B, C and D). Cells were stained with monoclonal mouse anti-Ty (Fluorescein channel) and rabbit anti-Aldolase (Alexa 568 channel). Nucleus and kinetoplast are stained with DAPI. Anti-TY was used for detection of TbLysoPLA.

### A peculiar PTS-1 signal allows dual localisation

The analysis of the primary sequence shows a C-terminal signal close to PTS1, “SKS” which is specific for this protein. We hypothesized that this PTS1-like signal could be important for the subcellular localisation of TbLysoPLA. The strategy was to express different mutant versions of TY-tagged TbLysoPLA in the ΔTbLysoPLA genetic background to study the distribution of the overexpressed proteins, upon biochemical fractionation and immunofluorescence analyses^33^. To validate our approach the TY-tagged TbLysoPLA protein expressed in the TbLysoPLA-null background showed the same distribution of TbLysoPLA as the native TbLysoPLA detected with the anti-LysoPLA antibodies in the wildtype context (Figure 4B, left panel). The upper band corresponds to the Ty-tagged protein and the lower band to the wild-type protein. This result is confirmed by hypotonic lysis fractionation (4B, middle) and by immunofluorescence analyses (4B, right). To determine the possible role of the PTS1-like SKS motif in the partial glycosomal localisation of TbLysoPLA, the SKS motif was removed (TbLysoPLA^noSKS_NTy^) or replaced by a canonical PTS1 motif (TbLysoPLA^SKL_NTy^) and the recombinant TY-tagged proteins expressed in the ΔTbLysoPLA background ((ΔTbLysoPLA-TbLysoPLA^SKS_NTy^ and (ΔTbLysoPLA-TbLysoPLA^SKL_NTy^ respectively). Digitonin treatment (Figure 4C, left), hypotonic lysis fractionation (Figure 4D, middle) and immunofluorescence (Figure 4C, right) show that TbLysoPLA without PTS1-like signal has the same cytosolic distribution as enolase, i.e. all TbLysoPLA localises in the cytoplasm. Thus, the SKS C-terminal signal is required for glycosomal targeting. In contrast, the TbLysoPLA^SKL_NTy^ protein is exclusively localised in the glycosomal compartment (Figure 4D). Together, these data showed that SKS PTS1 motif is responsible for the partial glycosomal localisation of LysoPLA.

### TbLysoPLA secreted by BSF and anti-TbLysoPLA antibodies are produced by Trypanosoma brucei gambiense infected mice

To confirm previous observations that TbLysoPLA in secreted by *Trypanosoma brucei gambiense* (*Tbg*) BSF^24^, we incubated the 427 *T. brucei* strain in a medium promoting the secretion / excretion of proteins and analysed the presence of TbLysoPLA in these fractions by Western Blotting^34^. As shown in Figure 5A (left), TbLysoPLA is well detected in the protein material excreted by the parasites. We could also reveal the enolase, which is a known excreted/secreted protein, but not threonine dehydrogenase which a soluble mitochondrial enzyme that should not be detected. In the figure 5A (right) we can appreciate the coomassie profile of the ESA compared to the whole cell extract.

**Figure 5:**
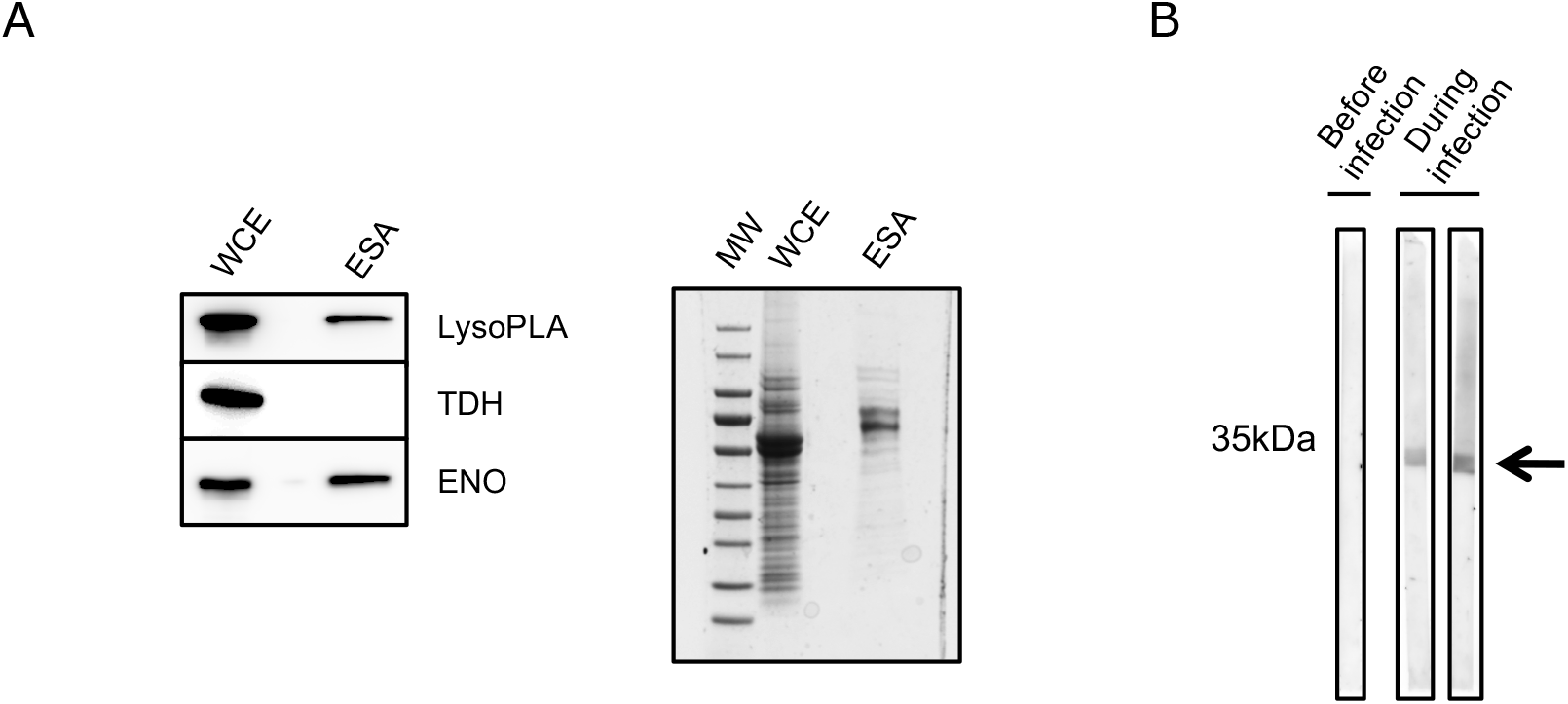
TbLysoPLA as a potent excreted/secreted factor. A / Left: Detection of LysoPLA in whole cell extract (WCE) of WT Tb BSF and in excreted/secreted material antigens (ESA). Specific anti-TbLysoPLA was used for detection, enolase was the positive control and TDH was used as a negative control. Right: Commassie stating showing WCE and ESA. B / Antibodies directed against TbLysoPLA are produced during infection Recombinant full-length TbLysoPLA was resolved by SPS-PAGE followed by a Western Blot. Mice serum taken before and during infection by *T.b gambiense* were used as primary antibody. Signal is detected only during infection (arrow).

The detection of TbLysoPLA in the excreted/secreted material released by the parasites, may favor its recognition by the immune system of the infected host. To test if specific antibodies are raised against TbLysoPLA during trypanosome infection, we tested the sera of healthy mice and experimentally infected mice with *T.b gambiense* (*Tbg*)^35^. The recombinant protein expressed in *E. coli* was resolved by SDS-PAGE and these sera were used for detection by Western Blotting as previously described by others^13^. As shown in Figure 5B, a signal is only obtained for samples collected during infection, indicating that infected mice develop an immune response against TbLysoPLA.

## Discussion

Only one phospholipase A1 had been previously characterized in African trypanosomes^18,19^.This enzyme is cytosolic, not excreted and involved in the synthesis of *lyso*-PC metabolites. In this study we describe TbLysoPLA, a new phospholipase with *an in vitro* A1 activity that showed a dual cytosolic and glycosomal localisation. The glycosomal localisation is due to a unique non-canonical PTS1 signal (SKS). In addition, the enzyme is found in the material excreted by the parasites and antibodies against this protein were detected in mice experimentally infected with *T.b gambiense*. We found that TbLysoPLA is not essential for the survival of parasites cultured under standard *in vitro* conditions, nor during an infection in mice. Indeed, we have seen using two reverse genetics approaches (i.e. RNA interference and gene knock-out), that parasites with no TbLysoPLA expression grow at the same rate as the parental strain. In addition, the KO cell-line is as virulent as the parental cell-line in a virulent mouse model of infection. In the future, it would be interesting to test the chronic effect of the absence of TbLysoPLA using a pleomorphic strain that cause longer infection^36^. Nevertheless, from our results we can conclude that TbLysoPLA is not a critical virulence factor for the establishment of an *in vivo* infection.

Among African trypanosomes, T. *brucei* has the highest PLA1 activity^15^, in particular compared to *T. congolense* for which it is very low. Interestingly, PLA1 activity was detected in the blood plasma of rabbits infected with *T. brucei*^17^. Thus, one may consider that this activity helps the parasites to penetrate the endothelium and other barriers, since *T. brucei* invades tissues while *T. congolense* remains vascular^37^. Another important function could be the detoxification of the environment, especially lysophospholipases (LPLs). We and others (our unpublished work,^38^) have observed that BSF do not metabolize phosphatidylcholine, but lysophospholipids decrease very strongly. TbLysoPLA could, as observed for most phospholipases A1^4^, also have a LysoPLA activity. Moreover it is interesting to notice that the gene is annotated as LysoPL. Given the versatility and diversity of sequences, it is complicated to classify these enzymes on its amino sequence, most can cleave several substrates: PL, LysoPL, di/tri glycerides. *In vitro* studies may not reflect the reality *in vivo*, and that is the reason why we have not focused our work on the specificity of substrates very precisely. This may be done later with the recombinant protein, in particular because there could be applications for the industry. To date, few PLA1 proteins have been expressed and purified. This type of protein could have a particular utility in the production of lipids that are used in cosmetics and in the food industry^10,39^.

TbLysoPLA is the second enzyme with an *in vitro* phospholipase A1 activity experimentally described in African trypanosomes. Moreover, it is the first enzyme displaying a phospholipase A2 activity *in vitro* ever described in trypanosomes. Its distribution is very peculiar since conventionally phospholipases are highly compartmentalized enzymes, in particular they are very often found associated with membranes where they play an important role for the remodeling of membrane lipids^3,10^. The first phospholipase A1 described in *T. brucei* PLA1a^19^ is located only in the cytosol and is not excreted, so it is unlikely that these two enzymes are functionally redundant since they do not show the same compartmentalisation. Like other protozoan parasites, African trypanosomes have several phospholipases genes in their genome in addition to PLA1 and LysoPLA (our unpublished work,^11,40^). An explanation for this expansion could be a possible functional redundancy. It will be necessary to study these proteins in greater detail in order to have a more complete and exhaustive understanding of the PLs function in trypanosomes.

This protein might not be distributed in the same way in the different kinetoplastids because not all of them have the PTS1 (Figure 1). Moreover in the genome of *T. brucei* there are only three proteins with the non-canonical PTS1 signal “SKS” and among them, only TbLysoPLA is addressed partially in glycosomes. In the absence of knowledge about the intracellular function of this protein, an explanation for this observation is difficult to advance. Nevertheless, we can speculate that this non-canonical signal is not effectively supported by the cytosolic chaperones responsible for transporting proteins to glycosomes. This has already been shown for other proteins in other cell types. As an example, the catalase of the yeast *H. polymorpha* has a non-canonical “SKI” signal^41^. A study showed that, by replacing this SKI sequence with a SKL sequence, the protein goes entirely into glycosomes^41^. However a part would form aggregates in glycosomes in which the protein would not be active. The import into the glycosomes would be so effective that it would not allow the protein to take its correct conformation. Thus the presence of a weak, less well-supported signal would leave time for the protein to take its correct folding. This could also be the case for TbLysoPLA. In this scenario, the main function of the protein could reside in the glycosomes and the cytosolic fraction would be largely unfolded and therefore not active. We could also consider that the cytosolic part is active. An excess in the cytosol can be deleterious for the cell because PL or LysoPL are toxins. A way to detoxify would be to excrete the enzyme outside of the cell, which is what we observe (figure 5).

Also, it is difficult based on our results to assert that LysoPL has a true PLA2 activity *in vitro*. For the moment no PLA2 has been purified or characterized so far^11^. Our study demonstrates that future investigation is needed to clarify this important question.

Finally, at some points TbLysoPLA is facing the serum of the animals either because it is released upon parasite destruction or because it is excreted/secreted as we suggested in this study. The enzyme is immunogenic enough that specific antibodies are raised upon infection. TbLysoPLA could be a very interesting target for the diagnosis of this parasitosis that is still not accurate specially for animals^42^.

The phospholipases of African trypanosomes are very little known, few genes are described and our study contributes modestly to fill this gap and pave the way for more studies.

## Materials and Methods

### Trypanosome growth and transfection

The bloodstream form of *T. brucei* Lister 427 90-13 (TetR-HYG T7RNAPOL-NEO), a Lister 427 221a line (MiTat 1.2) designed for the conditional expression of genes was cultured at 37°C in HMI-9 (*Iscove’s Modified Dulbecco’s Medium, Life Technologies* supplemented with 10% (v/v) heat-inactivated fetal calf serum, 0.25 mM β-mercaptoethanol, 36 mM NaHCO_3_, 1 mM hypoxanthine, 0.16 mM thymidine, 1 mM sodium pyruvate, 0.05 mM bathocuprone and 2 mM L-cysteine). Transfections were performed using Amaxa nucleofection method as previously described^43^.

### Production of recombinant TbLysoPLA and specific antibodies

A recombinant *T. brucei* full length LysoPLA fused to a GST tag on its N-terminal extremity, was expressed in *E. coli* One Shot BL21star (DE3) (Thermofisher) using the pGEX4T1 expression vector (GE Healthcare). Protein expression was induced at 37°C for 3 h using 0.5 mM isopropyl-D-thiogalactopyranoside. The cells were harvested by centrifugation, resuspended in PBS and sonicated. Proteins released in the soluble form were purified using Glutathione Sepharose 4B according to the manufacturer’s instructions (GE Healthcare). On-column thrombin digestion was performed to release the protein without the GST tag, then dialysed against PBS. Purified recombinant TbLysoPLA was used as an antigen to raise polyclonal antibodies. Two rabbits were injected 4 times at 15-days intervals using Covalab facilities (www.covalab.com).

### Site-directed mutagenesis

To mutate the serine 171 to alanine two complimentary primers were synthesized (supp table S3). The vector pGEX4T1/LysoPLA was used as a template and the amplification was performed using the *Pfu* ultra (Agilent, 600380). The PCR product was digested with *dpn1*, and then transformed in *E coli* XL1-blue. Plasmids were then extracted and sequenced (Eurofins genomics facilities) to confirm the presence of the mutation.

### Fluorometric phospholipase A assays

PLA1 activity was monitored using EnzCheck Phospholipase Assay Kits (Invitrogen, Life Technologies) according to the manufacturer’s instructions and as previously described in ^44^. Reactions were performed in black 96-well microplate for one hour at room temperature. The substrate PED-A1 is a Bodipy FL dye labelled phosphatidylethanolamine, the emission of which is dequenched upon PLA1 hydrolysis. Activities (30 μg per protein) were monitored by measuring fluorescence intensities at 485 nm excitation and 530 nm emission with an Optima microplate reader (BMG Labtech, Germany).

### Substrate specificity assay of recombinant TbLysoPLA

Two mixture of lipids were prepared by mixing their chloroform/methanol stocks (20 nmoles each) in a glass vial and drying on a nitrogen line, after which 50 ul of N-octyl-gucopyranoside (0.3% w/v) was added and sonicated. (Lipid mix 1: *lyso*-PC C17:0; PC (diC16:0); PC O-C16, 20:5PC (diC18:0); PC (18:0, 20:5).

Lipid mix 2, PA (diC16:0); PG (di C14:0); PS (di C14:0); PE (di C16:0); PE (O-C18:1, C18:1); PI (18:0, 20:4).

To these lipid mixed micelles/ vesicles was added 200ul of buffer 100 mM HEPES Na (pH 7.4), 10 mM MgCl_2_ and 1mM DTT ± 100ug of enzyme. This was sonicated again for a further 10 minutes prior to incubation overnight at 37C. The reaction was quenched by the addition of 750ul MeOH:CHCl_3_ (2:1) and the addition of an internal standard of either 25 nmoles PC (diC14:0) or 25 nmoles PA (diC17:0) to lipid mix 1 or 2 respectively. After vortexing for 20 minutes, chloroform and water were added to make biphasic and the lower chloroform rich layer was removed and dried down and stored at 4C in glass vial ready for mass spec analysis. Samples were suspended in MeOH:CHCl_3_ (2:1) and analysed on an Orbitrap mass spectrometer in both positive and negative mode

### Western blotting analyses

For Western blot analysis, total protein lysates of *T.brucei* BSF were separated by SDS-PAGE (4-20% Mini PROTEAN TGX stain-free precast gradient gels, Bio-Rad) and blotted on PVDF filters (Bio-Rad). The membranes were blocked with PBS 5% milk powder for 1 h at RT. Primary and secondary antibodies were diluted in PBS with 0.05% Tween 20 and 5% milk powder: rabbit anti-LysoPLA 1:1000 mouse anti-TY 1: 500 anti-aldolase 1:10000 anti-enolase 1:100000 Rabbit anti-PFR 1:10000 anti-mouse conjugated to horseradish peroxidase (KPL) 1:5000; or anti-rabbit conjugated to horseradish peroxidase (KPL) 1:10000. Revelations were done using Clarity Western ECL Substrate (Bio-Rad) according to the manufacturer’s instructions, pictures were acquired using a LAS4000 imager (GE Healthcare).

### Immunofluorescence Assay

Parasites grown in culture were collected by centrifugation, washed and fixed in paraformaldehyde as described elsewhere^37^. Slides were incubated with primary antibodies followed by Alexa Fluor 488-conjugated goat anti-mouse secondary antibody or Alexa Fluor 594-conjugated goat anti-rabbit secondary antibody (diluted 1:400) (Invitrogen). The nuclei were stained with DAPI (10 μg/mL) and cells were observed using a Zeiss Axio imager Z1 microscope; images were captured using Metamorph software (Molecular Devices). Images were processed using ImageJ software.

### RNA interference and gene knock-out

The inhibition by RNAi of the expression of the *TbLysoPLA* gene in the 427 BSF was performed by expression of stem-loop « sens/antisens » RNA molecules of the targeted sequences introduced into the pLew100 as previously described^31^. The sequence corresponding to the first 400 bp of the coding sequence was targeted. The sense and antisens fragments separated by 50 bp were cloned into the *Hind*III and *Xho*I restriction sites of the pLew100 vector.

Replacement of the *TbLysoPLA* gene by the blasticidin and puromycin resistance markers *via* homologous recombination was performed using DNA fragments containing a resistance marker gene flanked by the TbLysoPLA UTR sequences. The pGEMt plasmid was used to clone an *Hpa*I DNA fragment containing the blasticidin and the puromycin resistance marker gene preceded by the TbLysoPLA 5’UTR fragment and followed by the 3’ UTR fragment. Correct homologous integration of the resistance markers in the resulting drug resistant clones was analysed by PCR (see Fig. S4).

### Digitonin Permeabilisation

Trypanosomes were washed 2 times in cold PBS and resuspended at 6.5 × 10^8^ cells per mL (corresponding to 3.3 mg of protein/mL) in STE buffer (250 mM sucrose, 25 mM Tris, pH 7.4, and 1 mM EDTA) supplemented with 150 mM NaCl and the Complete™ Mini EDTA-free protease inhibitor mixture (Roche Applied Bioscience) and 1mM DTT. Cell aliquots (200 μL) were incubated with increasing quantities of digitonin for 4 min at 25°C, before centrifugation at 14,000 g for 2 min. Samples were then analysed by Western blot.

### Cell Fractionation

10^8^ parasites were washed in PBS and incubated in hypotonic lysis buffer (5 mM Na_2_HPO_4_, 0,3 mM KH_2_PO_4_) for 30 minutes at 4°C before centrifugation at 14.000g for 15 minutes. Material in the pellet was solubilised in SDS. Both pellet and supernatant were prepared for SDS-PAGE by adding Laemmli buffer^45^.

### Secretome of bloodstream form Trypanosomes

10^8^ parasites were washed in trypanosome dilution buffer (TDB, 20 mM Na_2_HPO_4_, 80 mM NaCl, 5mM KCl, 1 mM MgSO_4_, 20 mM glucose, pH7.4) and incubated in 15 mL of 50% serum-free HMI9, 50% TDB. During the experiment, cell viability was checked by microscopy. After 3 hours, cells were removed by centrifugation and Trypanosome-free medium was carefully taken, passed through a 0.22 μm syringe filter and concentrated 70 times using a Protein Concentrator 10,000 molecular weight cut-off filter unit (Pierce). Samples were then analysed by western blotting.

### Mice infection

*In vivo* experiments were performed with 10-weeks old male C57BL/6J mice, from Charles River Laboratories International. Mice were housed in a Specific-Pathogen-Free barrier facility, at Instituto de Medicina Molecular. The facility has standard laboratory conditions: 21 to 22°C ambient temperature and 12h light/12h dark cycle. Chow and water were available *ad libitum*. Animal experimentation work was performed according to EU regulations and approved by the Animal Care and Ethical Committee of Instituto de Medicina Molecular (AWB_2016_07_LF_Tropism).

The inoculum was prepared from thawed *T. brucei* cryostabilates and parasite motility was checked under an optic microscope. Mice were infected by intraperitoneal (i.p.) injection of 2000 parasites. At day 5 post-infection, animals were euthanized by CO_2_ narcosis and immediately perfused transcardially with pre-warmed heparinised saline (50mL phosphate buffered saline (PBS) with 250 μL of 5000 I.U./mL heparin). Organs were collected and snap frozen in liquid nitrogen.

### Parasite quantification in blood and organs

For parasitemia quantification, blood samples were taken daily from the tail vein and parasites counted manually in a hemocytometer (detection limit is 3.75 × 10^5^ parasites per mL of blood). Quantification of parasites in organs was performed by quantitative PCR of genomic DNA, as previously described in^36^.

## Supplemental

**Figure S1:**
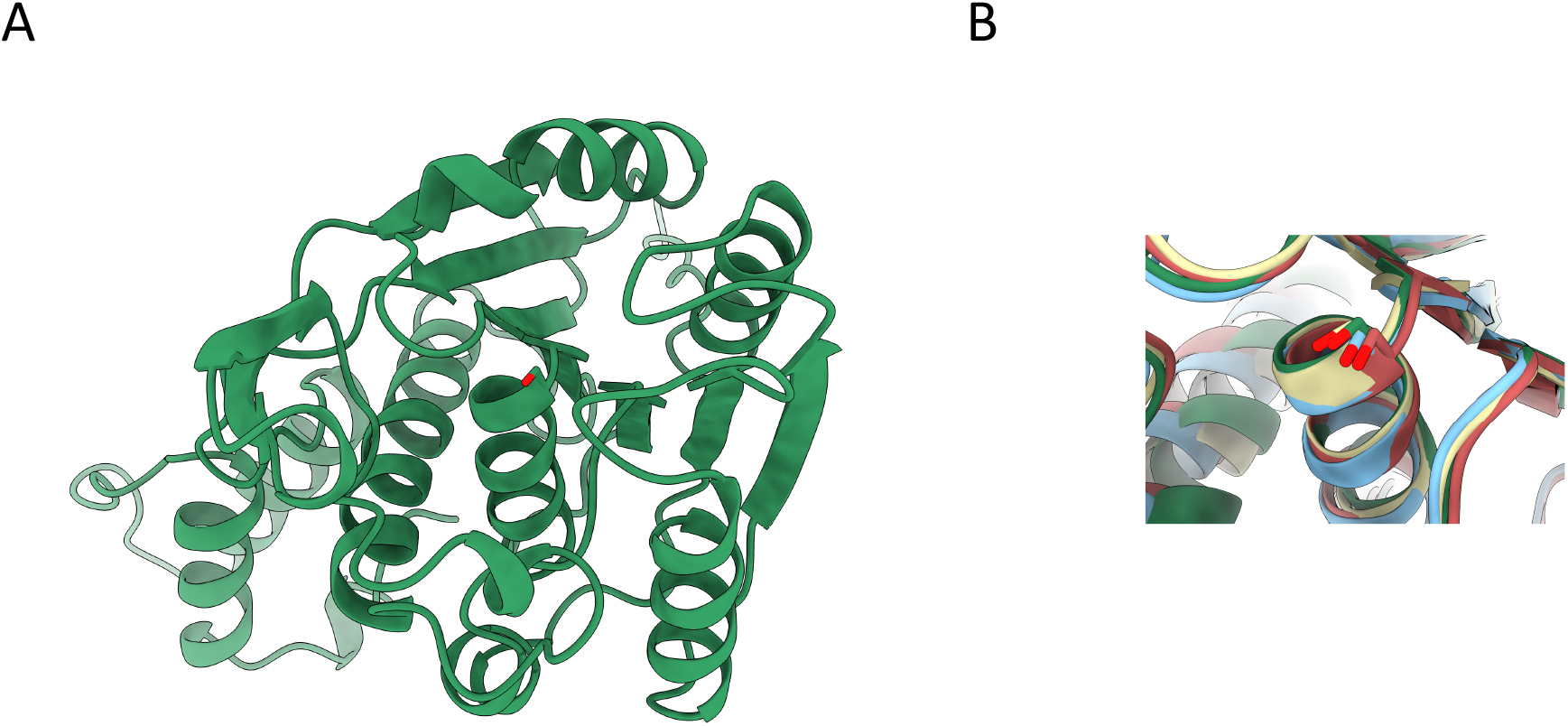
Structure prediction of LysoPLA using Alpha-Fold. A/ Prediction of TcLysoPLA using Alpha-fold. B/ Alignment of predicted catalytic site Pale cyan = Rhodobacter sphaeroides esterase (RspE) pdb id = 4FHZ; Pale Yellow = Francisella tularensis carboxylesterase (FTT258) pdb id = 4F21; Salmon = Human LYPLAL1= pdb id 3U0V; Pale blue = Human lysophospholipase A2 (LYPLA2) pdb = 6BJE.

**Figure S2:**
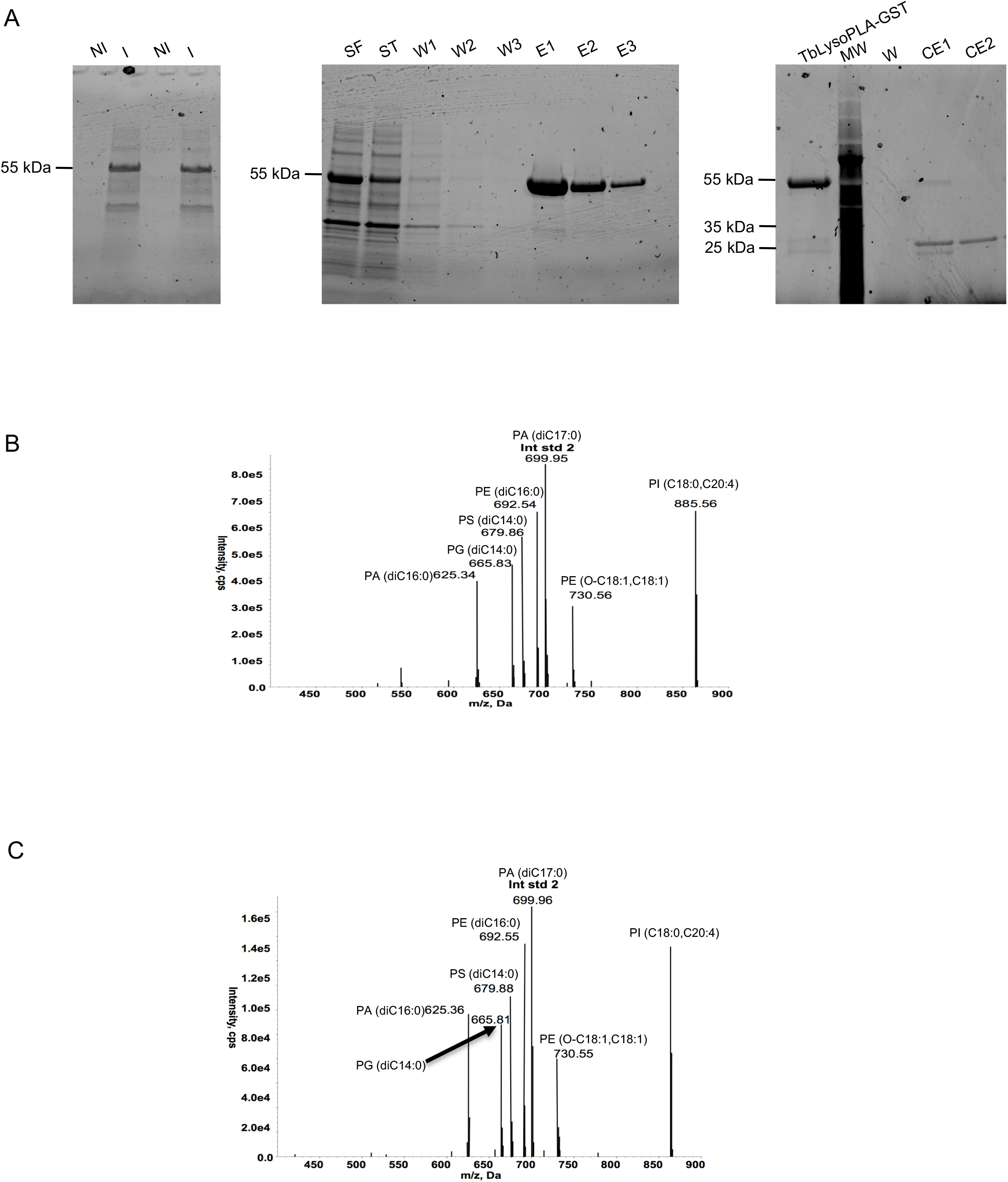
Expression and purification of TbLysoPLA. **A/** Left: Expression in *E. coli* BL21Star, non-induced, NI; induced with IPTG, I. Middle: Purification steps using Glutathion sepharose beads followed by glutathione elutions. SF (Soluble Fraction), FT (Flow Through), W1-2 (Washs), E1-3 (elutions with glutathion). Right: Thrombin clivage. MW, Molecular weight; W (Wash); CE1-2 (Clivage-Elution). B/ Substrate specificity of recombinant TbLysoPLA. TbLysoPLA was incubated (B) or not (C) with a Lipid Mix containing PA (diC16:0), PA (diC17:0) PG (diC14:0), PE (O-C18:1, C18:1), PE (diC16:0), PS (diC14:0), PI (C18:0, C20:4).

**Figure S3:**
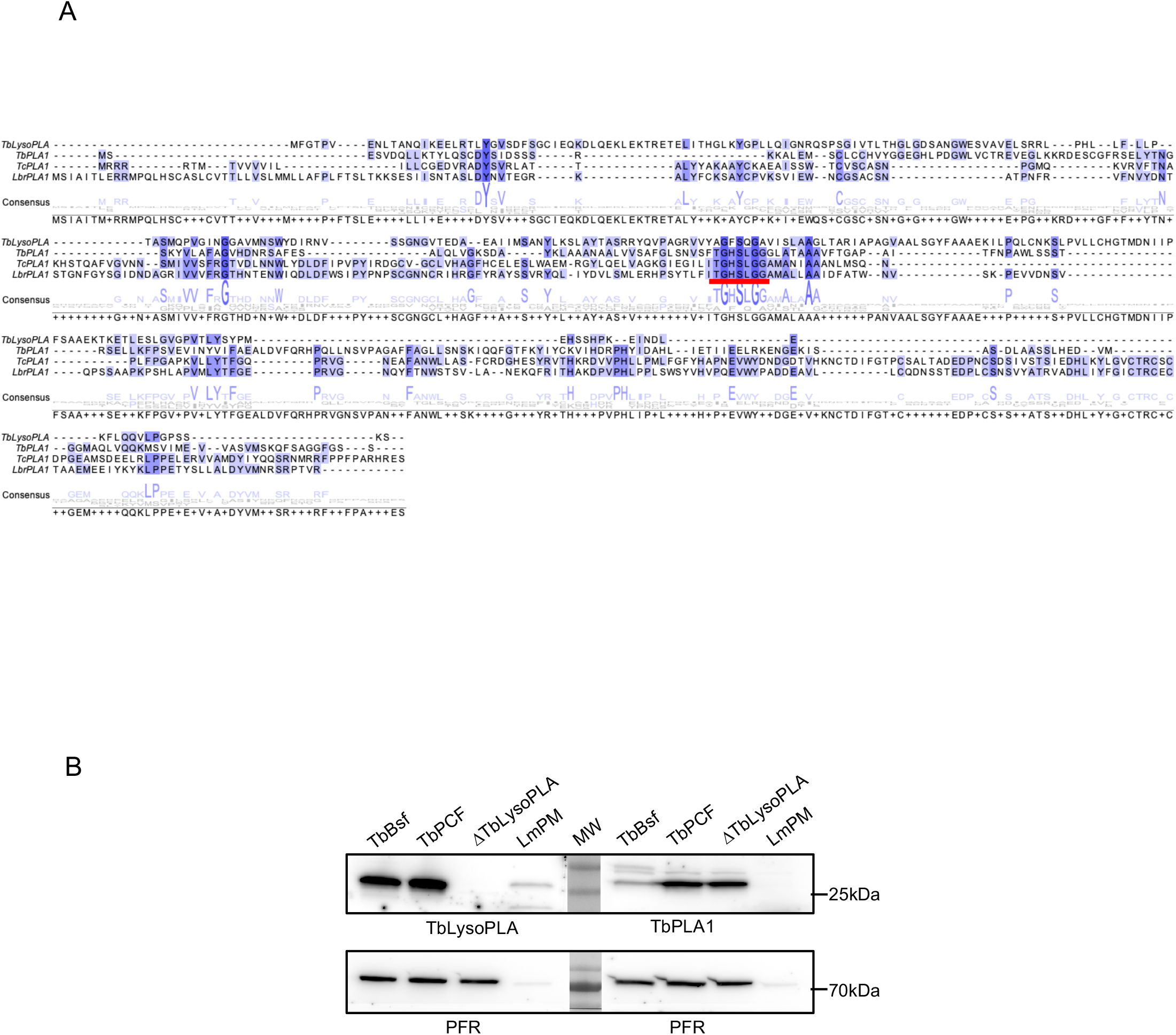
Comparison of TbLysoPLA and TbPLA1. A/ Multiple sequence alignment of TbLysoPLA and Tb, Tcr and Lbr PLA1. Amino acid sequences were aligned with Clustal Omega using basic settings and edited using the Jalview software. Dark blue contains conserved residues, white to light blue contains conservative changes. The lipase consensus pattern is underscored by a red lign. TbLysoPLA, *Trypanosoma brucei* LysoPLA (Accession number in GeneDB Tb927.8.6390); TbPLA1, *Trypanosoma brucei* PLA1 (Accession number CAG29794, ^19^); TcPLA1, *Trypanosoma cruzi* PLA1 (Accession number JN975637, ^13^); LbrPLA1, *Leishmania braziliensis* PLA1 (Accession number KJ957826, ^14^). B/ Western Blot analysis using anti-TbLysoPLA and anti TbPLA1b antibodies. Total protein extracts were resolved by SDS-PAGE and transferred on nitrocellulose as described in materiel and method section. Membranes were probed with anti-TbLysoPLA, anti-TbPLA1and anti-PFR. TbBsf, *T. brucei* Bloodsream form; TbPCF, *T. brucei* Procyclic form; TbLysoPLA; TbLysoPLA knock-out cell-line Tb bloodstream form; LmxPM, *Leishmania Mexicana* promastigote form. PFR was used as loading control.

**Figure S4:**
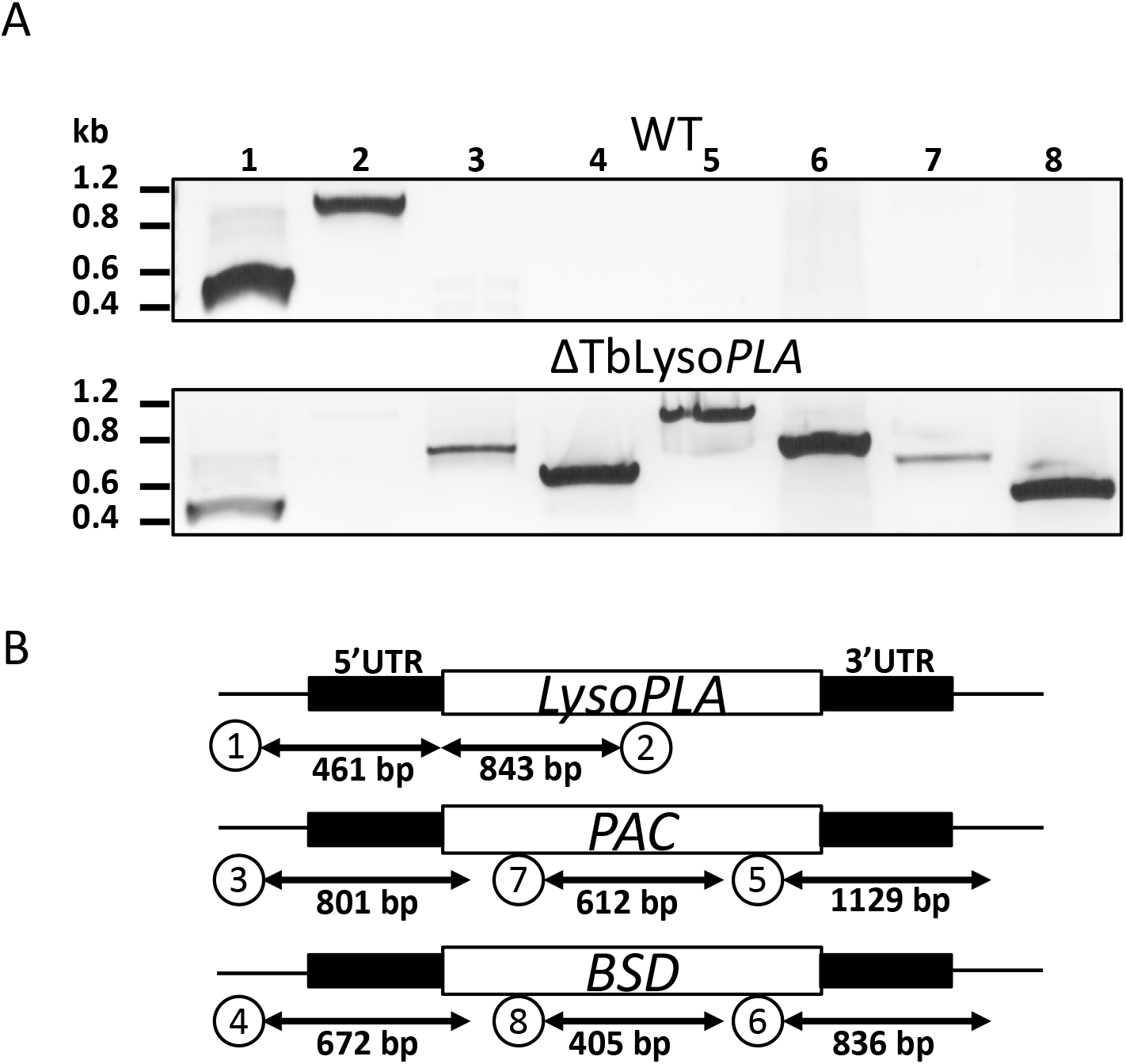
Generation of the ΔTbLysoPLA cell-line. TbLysoPLA alleles were replaced in the Tb427 parental cell-line by blasticidin and puromycin resistance genes. A / Representation of the wild type and recombinant loci and the PCR strategy to identify marker integration and wild-type loci. B / Analysis of genomic DNA extraction from parental cell-line (WT) or from a clone ΔTbLysoPLA by PCR amplification of the fragments presented in A (for primers see supplementary table).

**Table S1:** LysoPLA: percentage of identity among kinetoplastids. Protein sequences were aligned using Clustal Omega Algorythm with basic settings then identity matrix was retrieved.

|  | TbLysoPLA | TbgLysoPLA | TevLysoPLA | TcoLysoPLA | TviLysoPLA | TcrLysoPLA | LmLysoPLA |
| --- | --- | --- | --- | --- | --- | --- | --- |
| TbLysoPLA | 100 |  |  |  |  |  |  |
| TbgLysoPLA | 99.64 | 100 |  |  |  |  |  |
| TevLysoPLA | 99.64 | 99.64 | 100 |  |  |  |  |
| TcoLysoPLA | 70.61 | 70.61 | 70.61 | 100 |  |  |  |
| TviLysoPLA | 52.17 | 52.17 | 52.17 | 51.19 | 100 |  |  |
| TcrLysoPLA | 62.86 | 62.50 | 62.50 | 63.44 | 54.33 | 100 |  |
| LmLysoPLA | 53.82 | 53.82 | 53.82 | 53.28 | 43.60 | 57.61 | 100 |

**Table S2:** Comparison of TbLysoPLA and TbPLA1. Protein sequences were aligned using Clustal Omega Algorythm with basic settings then identity matrix was retrieved.

|  | TbLysoPLA | TbPLA1a | TcrPLA1 | LbrPLA |
| --- | --- | --- | --- | --- |
| TblysoPLA | 100 |  |  |  |
| TbPLA1a | 16.06 | 100 |  |  |
| TcrPLA1 | 16.29 | 20.83 | 100 |  |
| LbrPLA | 18.27 | 17.25 | 37.50 | 100 |

**Table S3:**
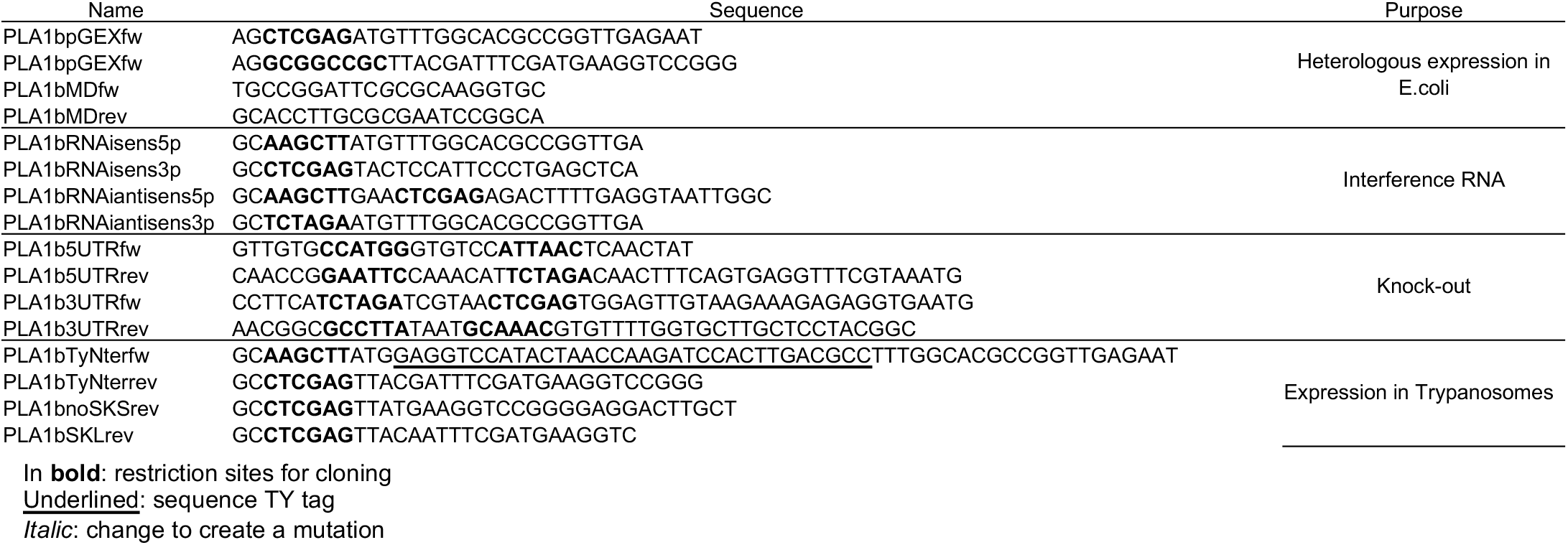
Primers used for PCR.

## Author Contributions

L.R and LM.F conceived the study; S.M, A.L, M.T, T.B-R, TK.S and L.R Conducted the experiments. S.M, A.L, M.T, T.B-R, LM.F, T.S and L.R analysed the data. S.M, M.T, T.B-R, LM.F, TK.S and L.R wrote the manuscript. S.M, M.T, T.B-R, F.B, LM.F, TK.S and L.R reviewed the manuscript. All authors approved the manuscript prior to submission.

## Supplementary informations

accompanies this paper.

## Ethics statement

All animal experimental work was performed in the Rodent Facility of Instituto de Medicina Molecular, which complies with Directive 2010/63/EU (transposed to the Portuguese legislation by Decreto-lei 113/2013) and follows the FELASA guidelines concerning laboratory animal husbandry and use. All animal research projects to be carried out at iMM are reviewed by the Animal Welfare Body (ORBEA-iMM) to ensure that the use of animals is carried out in accordance with legal requirements and following the 3R’s principle. The tasks involving animals were approved under the project AWB_2016_07_LF_Tropism with License number 018889\2016, issued by the local competent authority – Direcção Geral de Alimentação e Veterinária, being in agreement with the current legislation and with the recommendations for responsible use of animals. The ARRIVE guidelines were used for the reporting of in vivo experiments maximising the quality and reliability of published research, and enabling others to better evaluate and reproduce it.

## Acknowledgements

We are grateful to Dr Paul Lesbats for the help and discussions with structure prediction.

